# Synaptic and Somatic Targeting of ArcLight, a Genetically Encoded Voltage Indicator

**DOI:** 10.1101/2024.11.03.621404

**Authors:** Sheng Zhong, Lawrence B. Cohen

**Author notes:** Corresponding Author: Dr. Sheng Zhong, who currently work in Dept. of Molecular & Cellular Biosciences, University of Cincinnati College of Medicine, OH, USA.

## Abstract

Voltage signals in neurons are highly compartmentalized, which can influence their specific functions within neuronal circuits. Targeting of a genetically encoded voltage indicator (GEVI) to specific subcellular compartments can enhance the signal-to-noise ratio and provide more precise information about the location and timing of synaptic firing across different neuronal regions, reducing spatiotemporal signal convolution. To achieve subcellular targeting of the GEVI, ArcLight, we utilized five different postsynaptic targeting sequences (*Shaker* K^+^ channel C-terminus, stargazin C-terminus, rat Neuroligin-1 C-terminus, and anti-homer1 nanobodies HC20 & HC87) to direct ArcLight expression to the excitatory postsynaptic density. Additionally, we assessed a presynaptic-targeting tag (rat Neurexin-1β C-terminus) and a somatodendritic targeting tag (Kv2.1-Lk-Tlcn C-terminus). Patch clamp experiments in HEK293 cells showed that the targeting tags used in this study did not significantly alter ArcLight’s voltage sensitivity compared to controls. AAV infection in the mouse olfactory bulb demonstrated that the subcellular targeting sequences effectively localized GEVI expression to specific compartments of mitral/tufted cells, including postsynaptic densities, presynaptic terminals, and somatodendritic regions. Furthermore, i*n vivo* voltage imaging in mice expressing targeting-enhanced ArcLight variants revealed odorant-evoked responses similar to those observed with the original ArcLight. This indicates that subcellular targeting did not significantly impact the voltage sensing capability of ArcLight in mitral/tufted cells.

## INTRODUCTION

Genetically encoded voltage indicators (GEVIs) or calcium Indicators (GECIs) that expressed in specific cell types have emerged as powerful tools for advancing our understanding of brain function [1, 2]. While calcium signals provide valuable information, intracellular calcium dynamics are substantially slower in order of magnitude than voltage signals in neurons and are dynamically modulated by multiple cellular processes [3–5]. Thus, voltage measurements on neuronal signals are essential for understanding neuronal communication networks [6, 7]. Furthermore, neurons are highly spatially compartmentalized that engage in dynamic, spatially restricted signaling and information processing [8, 9]. To decipher neural circuits across networks of various scales, it is fundamentally crucial to understand how individual neurons integrate thousands of synaptic inputs and how axons filter and reshape somatic signals [10–12]. Targeting GEVIs to specific subcellular compartments within neurons could provide deeper insight into neural circuitry.

In General, subcellular targeting of GEVIs offers two major potential benefits. First, restricted expression can improve the signal-to-noise ratio by reducing background fluorescence from non-relevant regions. Second, subcellular compartment targeting of GEVIs can avoid signal convolution facilitating easier interpretation of recorded signals. Voltage signals in different neuronal compartments exhibit non-uniform dynamics during propagation [13]. Collecting signals from a whole neuron or a large area across different neuronal compartments can lead to spatiotemporally convoluted signals of GEVI fluorescence [14, 15]. By targeting GEVIs to specific functional compartments, more precise information about the location and timing of synaptic inputs or outputs can be obtained. Such spatiotemporally dynamic information is critical for interpreting neural communication network functions at the synaptic level and how different neuronal compartments interact and coordinate to process information flow. Technically, multiphoton imaging can achieve lateral resolutions of 1.8 um and axial resolution of 10 um [16], or scan up to 1500 µm deep in the brain with high frequency at cellular resolution [17–20]. The advances in multiphoton microscopy have opened possibilities for exploring the brain *in vivo* with higher spatiotemporal resolution [21].

Although most GEVIs are expressed uniformly over the cell membrane, the soma targeting expression of GEVIs has been reported [22–26]. Postsynaptic targeting constructs using different targeting sequences have also demonstrated effective for postsynaptic targeting of a GECI [27] or visualizing synapses [28, 29]. In this study we started with the GEVI, ArcLight [30], which has several advantages for cellular imaging, including good subcellular trafficking and cell membrane targeting, bright fluorescence and a relatively high voltage sensitivity [14, 31, 32]. Several targeting sequences were employed for targeting ArcLight to different functional compartments of the mitral/tufted neurons in mouse olfactory bulb. The rat Neurexin-1β C-terminus was employed for presynaptic targeting, and the Kv2.1-Lk-Tlcn tag was used for somatodendritic targeting. Five different postsynaptic targeting sequences were added at the C-terminal of ArcLight for dendritic expression, including the *Shaker* K^+^ channel C-terminus (SKC), the rat stargazin C-terminus with phosphomimetic charge mutations, the rat Neuroligin-1 C-terminus, and two anti-homer1 nanobodies HC20 & HC87. The details of these targeting sequences are given in the **methods** section. These targeting sequences act as sorting/trafficking signals and/or anchoring motifs that contribute to targeting ArcLight molecules to the presynaptic, somadendritic, or postsynaptic membrane and prevent ArcLight from diffusing away from targeted spots. We verified the voltage sensitivity of these ArcLight targeting constructs with patch clamp recording on HEK293 cells and found that these ArcLight constructs still kept robust voltage sensitivity compared with that of the original ArcLight [33].

The highly organized neuronal layers in the olfactory bulb facilitate the distinction of targeted GEVI expression in different compartments. The *in vivo* subcellular expression of these ArcLight targeting constructs in mouse olfactory bulb demonstrated the specific expression in different compartments of mitral/tufted cells, with varying specificity and efficiency. We verified the voltage response of targeted ArcLight using *in vivo* voltage fluorometry in the mouse olfactory bulb under odorant stimulation. The targeting-enhanced ArcLights variants showed similar odorant-evoked responses to the original ArcLight, indicating that the subcellular targeting did not significantly affect olfactory function. These results suggest that subcellular targeting of ArcLight to different functional compartments can be achieved and will be a useful method for improving our understanding of neural networks.

## METHODS

The original ArcLight-Q239 was modified with a Golgi trafficking signal sequence (Golgi-TS) and an endoplasmic reticulum (ER) export signal [34, 35] from the Kir2.1 potassium channel [36, 37] to enhance subcellular trafficking to the plasma membrane [31, 38, 39]. In addition, the trafficking-enhanced ArcLight was also codon-optimized for use in mammals [39]. All GEVI constructs used in this study were based on the optimized ArcLight (*Plasmid #100038, Addgene, Cambridge, MA*) derived from the original ArcLight-Q239 *(Plasmid #36856, Addgene, Cambridge, MA)*.

For the subcellular targeting of ArcLight to different compartments, different targeting sequences were tagged onto the C-terminal of ArcLight, respectively (*Figure 1*). The names of the ArcLight targeting constructs below consist of a prefix (“pre-”, “post-”, or “soma-”), indicating target compartment, followed by “targeting tags” as a suffix.

**Figure 1.**
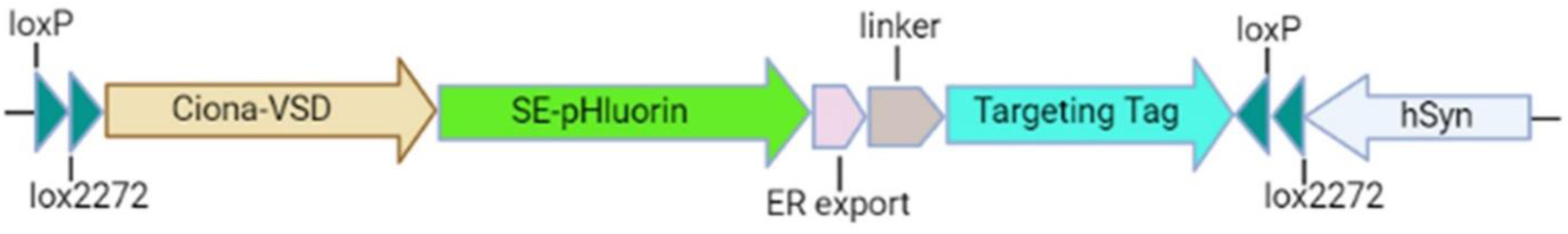
Schematic representation of AAV constructs for ArcLight targeting. The ArcLight targeting sequences were merged into a Cre-dependent expression cassette of AAV-DIO vector (**D**ouble-floxed **I**nverted **O**rientation with *LoxP* and *Lox2272*) with the human synapsin I promoter. SE-pHluorin represents Super-Eclipse pHluorin with A227D mutation.

### Presynaptic Targeting

#### preArcLight-rNxn

Neurexin-1β is a presynaptic adhesion protein, which interacts transsynaptically with neuroligin, critical for synaptic formation and function [40, 41]. It has been determined that the neurexin C-terminus dominates the subcellular trafficking and synaptic targeting of neurexin [42, 43]. Both pre-mGRASP [28] and pre-SynView [29] used the rat neurexin-1β for presynaptic targeting. We decided to employ the C-terminus of rat neurexin-1β (residues 415-468) for presynaptic targeting.

The presynaptic targeting sequence of rat Neurexin-1β tag is as follows: (KYRNRDEGSYHVDESRNYISNSAQSNGAVVKEKQPSSAKSANKNKKNKDKEYYV).

### Somatodendritic Targeting

#### somaArcLight-KvLkTn

The Kv2.1 potassium channel is predominantly clustered at the soma and proximal dendrites in mammalian brain neurons [44]. It was suggested that a 65-residue domain (residues 540-604) at the Kv2.1 C-terminus drives the post-Golgi transport vesicles for somatic targeting [22, 45–47]. Alternatively, the C-terminal sequence of telencephalin (Tlcn, residues 896–912) has been reported sufficient for targeting somatodendritic membrane in several types of neurons [26]. A hybrid motif, *Kv2.1C-linker-TlcnC*, had proved to be the best somatodendritic targeting motif (Addgene plasmid #114377) [48], which was used in our soma-targeting construct.

The somatodendritic targeting sequence of the **Kv2.1-Lk-Tlcn** C-terminal tag is as follows: (QSQPILNTKEMAPQSKPPEELEMSSMPSPVAPLPARTEGVID-MRSMSSIDSFISCATDFPEATRFGSGSGSGSGSAESPADGEVFAIQLTSS).

### Postsynaptic targeting

#### postArcLight-rNL1

Neuroligin-1 is a postsynaptic adhesion protein at excitatory synapses [49], which interacts with neurexin across the synaptic cleft. The postsynaptic targeting construct of post-mGRASP used the mouse neuroligin-1 skeleton [28], including the signal peptide, a juxtamembrane domain, the transmem-brane domain and the C-terminus, whereas post-SynView only adopted the C-terminus of rat neuroligin-1 for postsynaptic targeting [29]. We employed a reduced C-terminus of rat neuroligin-1 (residues 772-843) for postsynaptic targeting, containing the dendritic targeting motif (residues 772-803) and the PDZ-binding motif (-TTRV). The Neuroligin-1 C-terminal tag didn’t include the whole C-terminus (residues 719-843) as the critical region (residues 746-758) may not be necessary for dendritic targeting although it contributes to enhanced AMPAR current [50].

The postsynaptic-targeting sequence of rat Neuroligin-1 tag is as follows: (VVLRTACPPDYTLAMRRSPDDVPLMTPNTITMIPNTIPGIQPLHTFNTFT-GGQNNTLPHPHPHPHSHSTTRV).

#### postArcLight-SKC

The *Drosophila melanogaster Shaker* K^+^ channel C-terminus, containing a PDZ binding motif (-ETDV), can direct the postsynaptic localization of a GECI (SynapCam) for the local detection of calcium influx in *C. elegans* neuron [27, 51, 52]. We utilized the *Shaker* K^+^ channel C-terminus (SKC, residues 503-655) to test the targeting expression in mammalian neurons.

The sequence of *Shaker* K^+^ channel C-terminal tag is as follows: (TSCPYLPGTLVGQHMKKSSLSESSSDMMDLDDGVESTPGLTETHPGRSAVAPFL-GAQQQQQQPVASSLSMSIDKQLQHPLQQLTQTQLYQQQQQQQQQQQNGFKQQQQQTQQQLQQQQSHTINASAAAATSGSGSSGLTMRHNNALAVSIETDV).

#### postArcLight-Stgz

Stargazin is an auxiliary subunit of the AMPA receptor that promotes postsynaptic trafficking of the AMPA receptor by dynamic phosphorylation [53–55]. The C-terminal tail of stargazin contains 9 conserved serine residues as putative CaMKII/PKC phosphorylation sites [56–58] and a PDZ-binding motif at the very end of the C-terminus [56, 59]. We employed the rat stargazin C-terminus (residues 203-323) (Addgene plasmid # 80406) [60] with three phosphomimetic mutations (S239D, S240D, and S253D). For promoting interaction with deeper PSD95 [61], a 15-amino acid linker (GGGGS)_3_ was added between ArcLight and the stargazin C-terminus.

The postsynaptic-targeting sequence of stargazin tag is as follows: (DRHKQLRATARATDYLQASAITRIPSYRYRYQRRSRDDSRSTEPSHSRDADPVGVKGFNTLP-STEISMYTLSRDPLKAATTPTATYNSDRDNSFLQVHNCIQKDSKDSFHANTANRRTTPV).

#### postArcLight-HC20/HC87

Besides the scaffolding protein PSD95, Homer1 could be another candidate for postsynaptic targeting as it was shown to be evenly distributed within the postsynaptic density (PSD) of glutamatergic excitatory synapses [62]. Homer1 is a postsynaptic scaffolding protein in excitatory synapses, which occupies a layer 30– 100 nm from the postsynaptic membrane [63]. We employed two anti-homer1 nanobodies, **HC20** and **HC87** (Addgene plasmid #135220 & #135223) [64], as anchoring tags that would bind to Homer1 [65]. A 15-amino acid flexible linker (GGGGS)_3_ was added between ArcLight and the anti-Homer1 nanobodies to reach deeper Homer1 layer underneath the synaptic membrane.

The postsynaptic targeting sequence of **HC20** is as follows: (MAEVQLVESGG-GLVQAGGSLRLS-CAASGLTVNMWYTGWFHQAPGKEREFVAGISQSGVRTYVADSVKGRFTISRDNAKKTVYLQMNSLKPEDTGVYYCAGGTWESEVRRGVLSSRGQGTQVTVSSGQAGQ).

The postsynaptic targeting sequence of **HC87** is as follows: (MAQVKLQESGG-GLVQVGDSLRLSCAASGRPFSDYYMAWFRQAPGKEREFVAVIGWSGLTTY-YADSVKGRFTISRDNAKNTGYLQMNSLKLEDTAVYYCAADLRRRLSLTEFDYWGQGTQVTVSSGQAGQ).

All the ArcLight targeting constructs were first fused into the pEGFP-N1 vector to test the voltage sensitivity by whole-cell patch clamp fluorometry in HEK293 cells. Then the ArcLight targeting constructs were transferred into a Cre-dependent expression cassette of AAV-DIO vector (**D**ouble-floxed **I**nverted **O**rientation with *LoxP* and *Lox2272*) with the human synapsin I promoter (the backbone is the same as the *Addgene #50457* plasmid) for making Adeno-associated viral (AAV) viruses. The AAV targeting constructs were packed into AAV capsids by the Penn Vector Core (Philadelphia, PA).

### Mammalian Cell Culture

HEK 293 cells (ATCC, Manassas, VA) were maintained in Dulbecco’s modified Eagle’s medium (DMEM, high glucose; Invitrogen, Carlsbad, CA) supplemented with 10% fetal bovine serum (FBS; Sigma-Aldrich, Saint Louis, MO), 1% penicillin (100 U/mL), and streptomycin (100 μg/mL) (Sigma-Aldrich). The HEK 293 cells were plated on 12 mm coverslips (EMS Acquisition Corp., Hatfield, PA) in a 24-well plate. Cell cultures were maintained in a humidified incubator at 37 °C in a 5% CO_2_ environment. Transient transfection of constructs was accomplished by using plasmid DNA (0.2 μg per 12 mm coverslip) and Lipofectamine 2000 (0.4 μL per well; Invitrogen).

### Whole Cell Patch Clamp of HEK293 cells

Patch clamp recordings were performed in a perfusion chamber with the bath temperature kept at 32−34 °C by a temperature controller (Warner Instruments, Hamden, CT). We used 3−5 MΩ glass patch pipettes (capillary tubing with 1.5/0.75 mm OD/ID-World Precision Instruments, Sarasota, FL) that were pulled on a P-97 Flaming/Brown type micropipette puller (Sutter Instrument Co., Novato, CA). The bath solution contained: 150 mM NaCl, 4 mM KCl, 2 mM CaCl_2_, 1 mM MgCl_2_, 5 mM D-glucose (Sigma-Aldrich, G5767), and 5 mM HEPES, pH 7.4, and was adjusted to 300 mOsm with D-Sucrose (Sigma, S0389). The pipette solution contained: 120 mM K-Aspartate (Sigma, A6558), 4 mM NaCl, 4 mM MgCl_2_, 1 mM CaCl2, 10 mM EGTA, 3 mM Na_2_ATP, 5 mM HEPES, pH 7.2 and adjusted to 300 mOsm with D-Sucrose or D-Glucose. Voltage-clamp recordings in the whole-cell configuration were performed using an Axopatch-200B amplifier (Molecular Devices LLC, San Jose, CA) with a holding potential of −70 mV.

### Animals and Viral Injection

All animal experiments and procedures, including the viral injection surgery and *in vivo* voltage imaging surgery, were performed in accordance with relevant guidelines and regulations, including the NIH *Guide for the Care and Use of Laboratory Animals* and a protocol approved by the Institutional Animal Care and Use Committee (IACUC) at Yale University. For all surgical procedures mice were anesthetized by intraperitoneal injection with a mixture of ketamine (90 mg kg^−1^, Ketaset, Zoetis Inc., Spain) and xylazine (10 mg kg^−1^, AnaSed, Akorn Inc., IL). Anesthesia was supplemented as needed to maintain areflexia, and anesthetic depth was monitored every 15 min by the pedal reflex. Atropine (0.2 mg kg^−1^, AtroJect SA, Henry Schein, OH) injected under the back skin and local anesthetic bupivacaine (8 mg kg^−1^, Hospira Inc., IL) injected under the skull skin were applied in all animal surgeries. Animal body temperature was maintained at approximately 37 °C with a heating pad underneath the animal body to maintain normothermia. For recovery manipulations, animals were maintained on the heating pad until awakening.

The AAV viruses were injected into the olfactory bulb of adult ***Tbx21-Cre*** mice (Jackson Laboratory, strain *#024507*) for targeted ArcLight expression in the mitral/tufted cells [66]. ***Tbx21-Cre*** mice of either sex between 3-6 months of age were used for AAV injection according to previously described procedures [67]. For AAV virus injections, anesthetized mice were administered subcutaneously with 0.1 mg kg^−1^ of buprenorphine (Par Pharmaceutical, NY) or 3.25 mg kg^−1^ of Ethiqa XR (Fidelis, NJ) under the back neck skin for analgesia. Petrolatum ophthalmic ointment (Dechra, KS) was applied to the eyes to prevent corneal drying. A small hole (< 300 μm) was carefully made in the skull directly above each olfactory bulb, and the AAV targeting viruses were intracranially injected into the olfactory bulb at a rate of 10 nL per 10 sec using a glass pipette and a microinjector (NanoLiter-2000, World Precision Instruments, FL). Following the AAV injection and incision suture, 14 days or more was allowed for the expression of ArcLight in the mouse olfactory bulb before *in vivo* voltage imaging of odorant response.

### AAV Virus Serotype

To infect mouse olfactory neurons, we made five AAV1 viruses with the targeting constructs of postArcLight-SKC, postArcLight-HC20, postArcLight-HC87, postArcLight-Stgz, and preArcLight-rNxn. We also made four AAV2 viruses with the targeting constructs of postArcLight-SKC, postArcLight-HC20, postArcLight-rNL1, and somaArcLight-KvLkTn. In general, AAV2 viruses resulted in less expression than AAV1 viruses in the mitral/tufted cells, whereas the targeting patterns did not change with different AAV serotypes (*Figure S1, S2*).

### Odorant Stimuli and Delivery

The odorant isoamyl acetate (isopentyl acetate, *Sigma-Aldrich 306967*) saturated vapor was used as the olfactory stimulus. The odorant was diluted from saturated vapor with cleaned air using a flow dilution olfactometer described previously [68]. The olfactometer was designed to provide a constant flow of air blown over the nares. The odorant was constantly injected into the olfactometer but sucked away via a vacuum that was switched off during odorant presentation.

### *In vivo* Voltage Imaging

*In vivo* voltage imaging was performed with the AAV-injected mice under anesthesia. The surgical procedure was similar to that of AAV injection. A wide-field epifluorescence microscope was used to image the odorant response in olfactory bulbs of AAV-injected mice. The bone above the olfactory bulb was carefully thinned for transparency under a stereomicroscope (Leica MZ16F). Then the exposed region was covered with PBS buffer and sealed with a glass coverslip. The anesthetized mouse was positioned in a stereotaxic head holder (KOPF Ins., CA) with a heating pad (37 °C) underneath the body to maintain normothermia. The dorsal surfaces of the olfactory bulbs were illuminated with a collimated modular Mic-LED Light Source (Prizmatix, Israel). We used a 488 nm excitation filter (BrightLine FF01-488/10-25, Semrock), a 506 nm dichroic beamsplitter (BrightLine FF506-Di03-25x36, Semrock), and a 496 nm long pass emission filter (Semrock FF01-496/LP-25). Images were acquired and digitized with a 256x256 pixel CCD camera (NeuroCCD-SM256, RedShirt Imaging LLC, GA) at a 125 Hz frame rate through a C-mount TV objective (5x F0.95, Computar, Japan). NeuroPlex software (RedShirt Imaging) was used to control the camera, display, and analysis of the fluorescence signals. The fluorescent signals were converted to ΔF/F by dividing the initial fluorescence by the mean of at least 200 ms before the odorant stimuli. Odor response values (ΔF/F) were calculated as the largest difference within a 2000 msec temporal window during the peak of the odor response and the time prior to odor stimulation.

### Histological Methods

In a subset of experiments after *in vivo* voltage imaging mice were given an overdose of euthasol (Virbac AH, Inc., TX) for euthanasia, followed by cardiac perfusion with PBS buffer and then 4% paraformaldehyde (Sigma-Aldrich 158127 or J.T. Baker S898-07). The brains including olfactory bulbs were dissected from the skull and left in 4% paraformaldehyde for a minimum of 3 days. Before cryosection each olfactory bulb was infiltrated in 30% sucrose overnight, embedded in optimal cutting temperature (O.C.T.) compound and frozen. The olfactory bulbs were cut in 30 μm thick coronal sections on a Leica CM3050S cryostat (Leica, Germany). Mounted sections were coverslipped with VECTASHIELD Mounting Medium with DAPI (Vector Labs, H-1500, CA). The fluorescence of the targeted ArcLight was measured directly using Zeiss confocal microscopes (Zeiss LSM-780 and LSM-900, Carl Zeiss Microsystems) with Zeiss objectives of Plan-Apochromat 20x / 0.8NA or 63x / 1.4NA DIC M27 Oil immersion. Confocal images were pseudo-colored using Zeiss Zen Blue (Carl Zeiss Microsystems) and cropped/combined with Adobe Photoshop/Illustrator (Adobe Systems Inc.). The ArcLight fluorescent intensity was measured using the Color-Split Channels and the Analyze-Histogram functions of ImageJ/FUJI software (NIH, USA).

## RESULTS

### Voltage Sensitivity in HEK293 cells

Whole-cell patch clamp fluorometry on HEK 293 cells was used to verify the voltage sensitivity of the ArcLight targeting constructs. We tested five constructs: three postsynaptic-targeting tags (postArcLight-Stgz, postArcLight-rNL1, and postArcLight-SKC), one with the presynaptic-targeting tag (preArcLight-rNxn) as well as one soma-targeting tag (somaArcLight-KvLkTn). The voltage responding traces of the ArcLight targeting constructs were shown side by side compared to the control ArcLight (*Figure 2*). The constructs of somaArcLight-KvLkTn and postArcLight-rNL1 exhibited reduced voltage sensitivity compared to the control ArcLight (*Figure 2A, 2C and 2F*). The on and off time constants of the constructs of preArcLight-rNxn, postArcLight-SKC, postArcLight-Stgz, and postArcLight-rNL1 were slightly slower than the control ArcLight (*Figure 2B, 2D, 2E, and 2F*). However, all targeting constructs kept the key properties as a GEVI. As HEK293 cell is functionally different with neurons and lack of synaptic structures for testing the specific targeting motifs, we need to verify the effectiveness of these targeting constructs in animal neurons.

**Figure 2.**
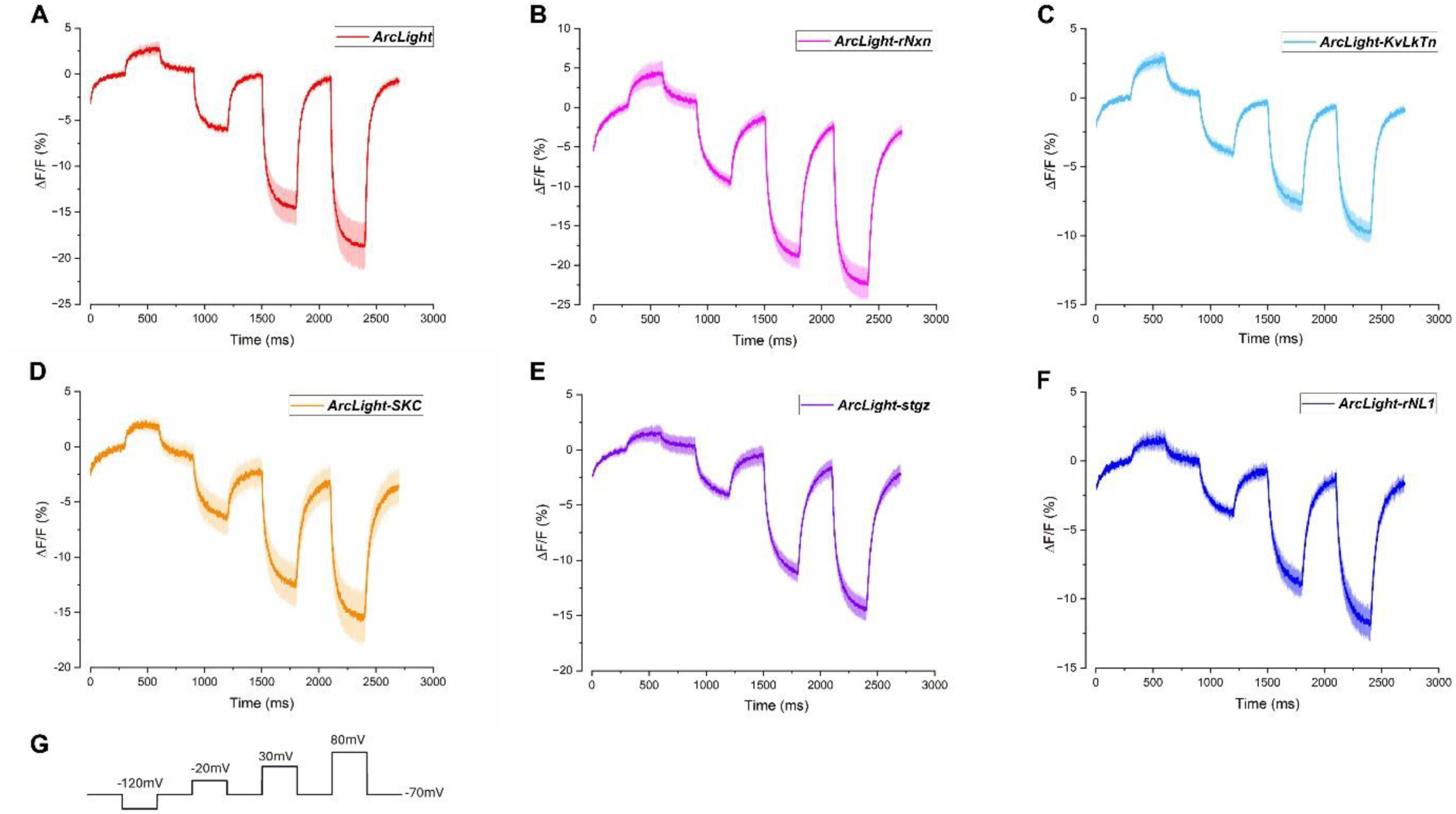
Representative voltage response fluorometry of the ArcLight targeting constructs measured by whole-cell patch clamp on HEK293 cells. (**A**) Averaged traces (the recorded cell number n=7) of the optimized ArcLight (*Addgene #100038*) as control. (**B-F**) The ArcLight targeting constructs with different targeting tags, including the presynaptic rat neurexin-1β (rNxn, n=7), the somatodendritic Kv2.1-linker-Tlcn (KvLkTn, n=8), and the postsynaptic *Shaker* K+ channel (SKC, n=5), stargazin (stgz, n=10), and rat neuroligin-1 (rNL1, n=9). (G) The voltage stimuli applied by patch clamp, aligned with the fluorescence intensity change of ArcLight constructs.

### Confocal Microscopy of the Expression of the Targeted ArcLights

Confocal microscopy revealed that the targeted ArcLight expression in mitral/tufted cells varied based on the targeting tags used. For example, preArcLight-rNxn was predominantly expressed in the internal plexform layer (IPL), while somaArcLight-KvLkTn was mainly expressed on the soma and proximal dendrites membrane in the the mitral cell layer (MCL) and external plexiform layer (EPL). Two postArcLight constructs (postArcLight-rNL1 and postArcLight-SKC) showed relatively higher expression in the glomerular dendritic tufts but exhibited distinct trafficking patterns. Detailed descriptions of the targeted ArcLight expression are provided below, with the AAV serotype indicated in parenthesis.

#### ArcLight (control, AAV1)

The control ArcLight without targeting sequence exhibited approximately uniformly across all compartment layers of the mitral/tufted cells, including the IPL. It was widely diffused on the membranes of the soma, dendrites, and axons. The ArcLight expression in the dorsal external plexiform layer (dEPL) was slightly higher than that in the MCL and ventral external plexiform layer (vEPL) (*Figure 3A*). Fluorescence intensity measurements on images from 8 animals confirmed a relatively even distribution of the control ArcLight (*Figure 5A*), with the dEPL showing somewhat higher expression.

**Figure 3.**
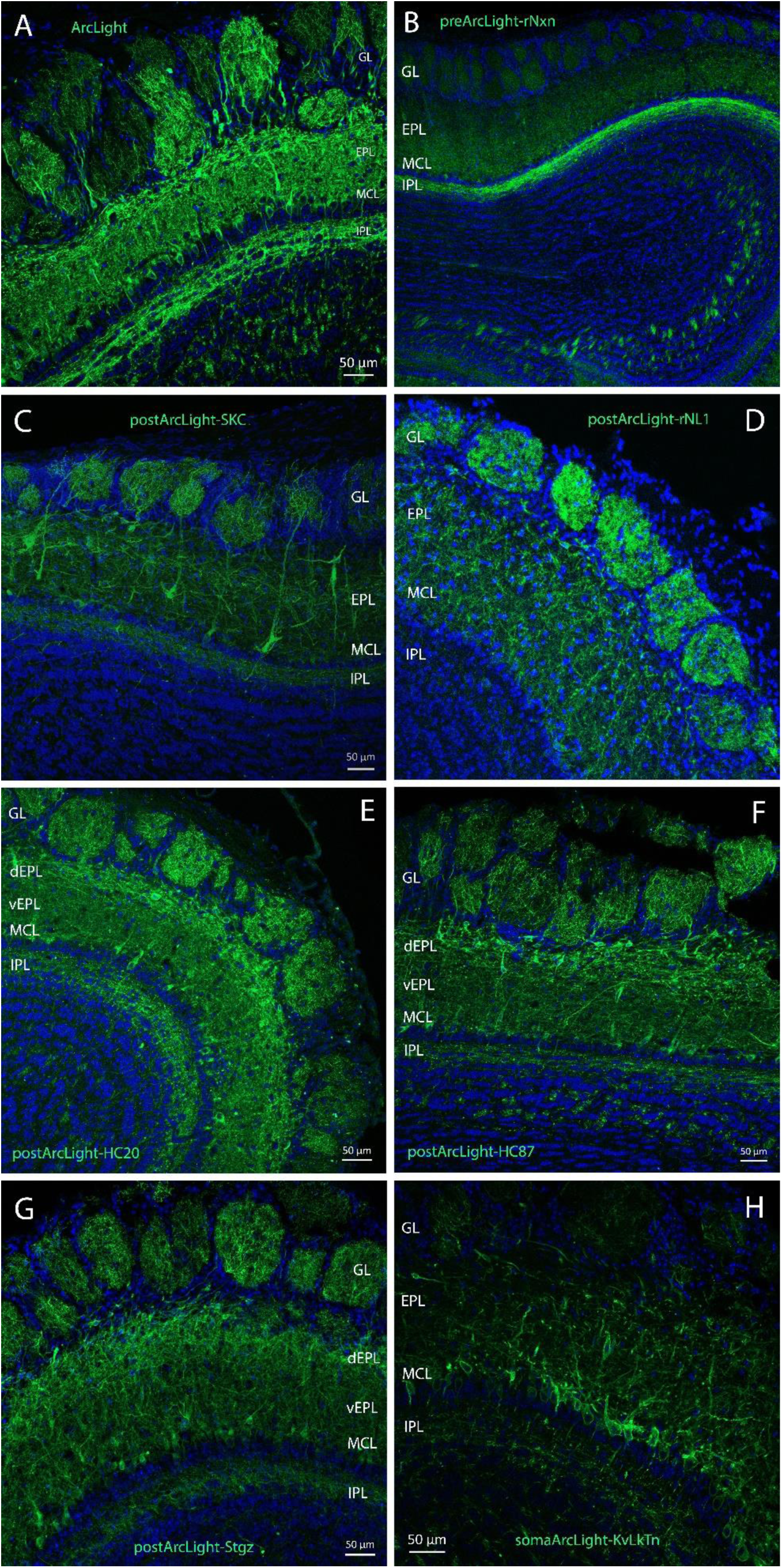
Confocal Microscopy of the Subcellular Expression of ArcLight targeting constructs in the mitral/tufted cells in the olfactory bulbs of *Tbx21-Cre* mice. **(A)** The control ArcLight expressed approximately evenly in every compartment of the mitral/tufted cells. **(B)** The preArcLight-rNxn expression was enriched in the internal plexiform layer (IPL). **(C)** postArcLight-SKC expressed mainly in the primary dendritic tuft at GL with minority expression in the lateral dendrites in EPL. postArcLight-SKC showed preferential distribution in the dendrite tuft in GL and lateral dendrite in the EPL and the soma membrane in MCL, while less than half the fluorescence was expressed in IPL. The SKC tag helps with cell surface targeting of ArcLight and showed some power of dendritic targeting, but some intracellular fluorescence also existed inside the soma and proximal dendrites. **(D)** postArcLight-rNL1 expressed mainly in the primary dendritic tuft at GL with minority expression in the lateral dendrites in EPL. **(E)** postArcLight-HC20. **(F)** postArcLight-HC87 expressed mainly in the distal EPL (dEPL) as well as the GL with limited expression in MCL and EPL. The targeted ArcLight expression was mainly distributed in the primary dendritic tufts in the GL and the lateral dendrites at the superficial EPL (sEPL) while the MCL and dEPL had dimer fluorescence. The IPL has limited expression. **(G)** postArcLight-Stgz expressed mainly in the distal EPL (dEPL) as well as the GL with limited expression in MCL and EPL. postArcLight-Stgz showed preferential distribution in different compartment of the mitral and tufted cells. The distal dendrite in the GL had higher expression than the lateral dendrite in the EPL and about 3 times higher than that of the IPL. Glutamatergic postsynaptic targeting construct with stargazin tag showed nonspecific distribution in the GL, EPL and IPL, maybe due to the phosphorylation situation of the stargazin C-terminus. **(H)** somaArcLight-KvLkTn expressed mainly in the soma and proximal dendrite membrane with limited expression in the primary and lateral dendritic membrane.

#### preArcLight-rNxn (AAV1)

The IPL comprises axon fibers from mitral/tufted cells in the olfactory bulb [69]. The presynaptic targeted ArcLight fluorescence of preArcLight-rNxn was predominantly distributed in the IPL while other layers appeared much dimmer (*Figure 3B*). The mean brightness in the IPL was 4 times larger than that in the GL (*Figure 4A*).

**Figure 4.**
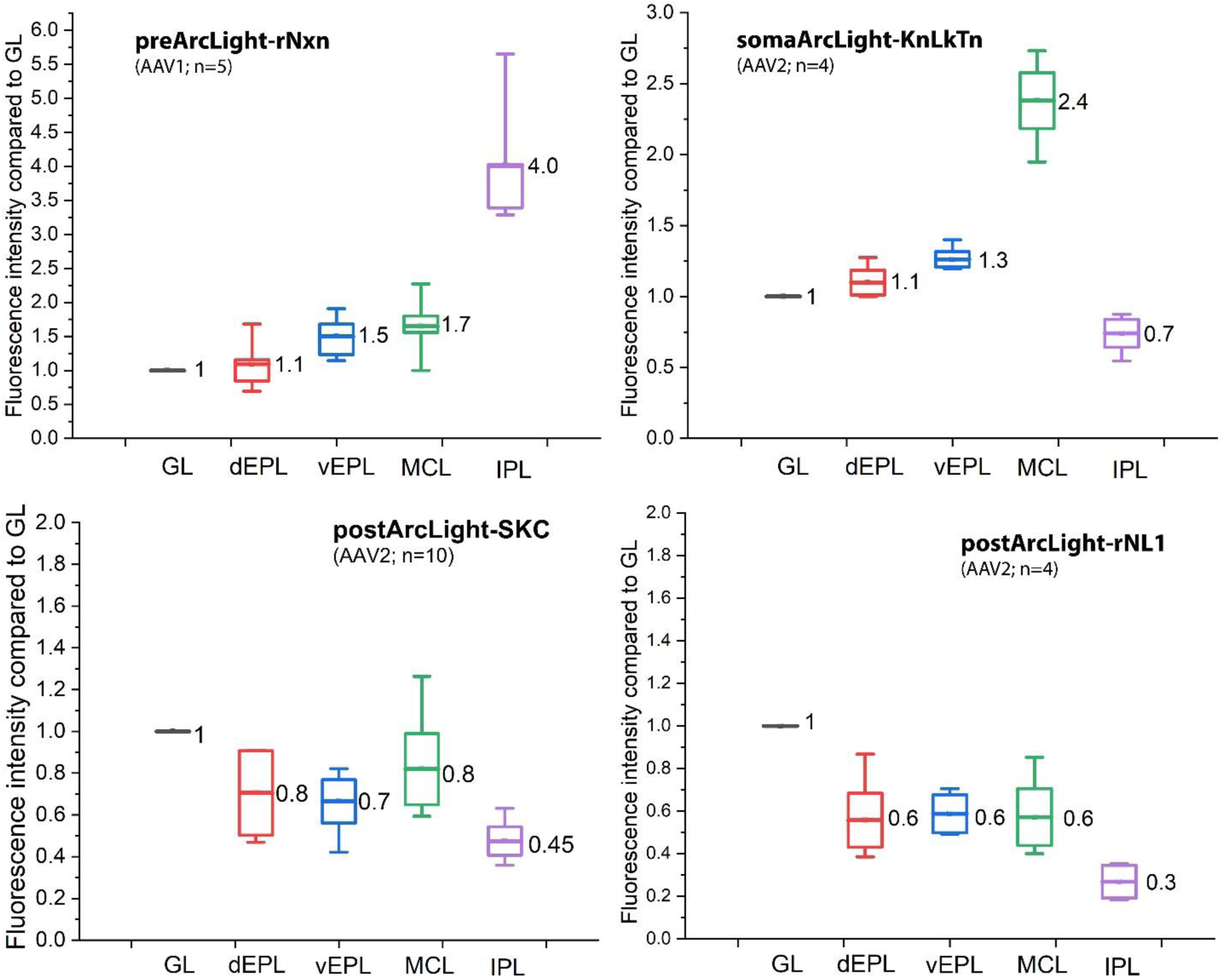
Fluorescence intensity measurements of ArcLight targeting constructs that showed specific subcellular expression. **(A) preArcLight-rNxn**. The brightest fluorescence was enriched in the internal plexform layer (IPL) which consists of axons from mitral/tufted cells. The mean brightness in IPL was 4.0 times as that in GL. **(B) somaArcLight-KvLkTn.** The fluorescence was enriched on the soma membrane and proximal dendrites of the mitral and tufted cells. The median value in MCL was 2.4 times as that in GL. **(C) postArcLight-SKC.** The fluorescence expression mainly distributed in the primary dendrites in the GL and in the lateral dendrites in the EPL and in the soma membrane in MCL. The IPL had an average fluorescence 2.2 times dimmer than that in the GL. **(D) postArcLight-rNL1**. The fluorescence expression was mainly distributed in the primary dendritic tufts in GL and with dimer fluorescence in the EPL and MCL, while average brightness in IPL was 3 times dimmer than that in the GL.

#### somaArcLight-KvLkTn (AAV2)

The soma-targeted ArcLight fluorescence was primarily expressed on the soma and proximal dendrites of mitral/tufted cells (*Figure 3H*). The primary dendritic tufts in the GL and lateral dendrites in the EPL showed minimal expression while the IPL exhibited even dimmer fluorescence. The mean brightness in MCL was 2.4 times greater than in the GL (*Figure 4B*).

#### postArcLight-SKC (AAV2)

The expression of the postsynaptic targeted ArcLight with SKC tag was unusually distributed along the primary dendrite axis from soma towards the dendritic tuft in the GL, while much dimmer fluorescence in the lateral dendrites in the EPL and in the soma membrane in MCL (*Figure 3C*). The mean brightness in the GL was more than two times brighter than in the IPL (*Figure 4C*). There were some intracellular aggregations of ArcLight fluorescence, implying a subcellular trafficking problem on cell membrane targeting. See the **discussion** section for more detail.

#### postArcLight-rNL1 (AAV2)

The expression of the postsynaptic-targeted ArcLight with rNL1 tag was preferentially distributed in the dendritic tuft in the GL and dimmer fluorescence on the lateral dendrites in the EPL, while minimal fluorescence in the IPL (*Figure 3D*). The mean brightness in the GL was three times brighter than in the IPL (*Figure 4D*).

#### postArcLight-HC20 (AAV1)

The postsynaptic-targeted ArcLight with anti-homer1 nanobody HC20 tag exhibited a slightly preferential distribution in dendritic tufts within the GL and EPL (*Figure 3E*). The mean brightness of ArcLight in the GL and EPL were 1.5∼1.8 times brighter than that in the IPL (*Figure 5C*).

**Figure 5.**
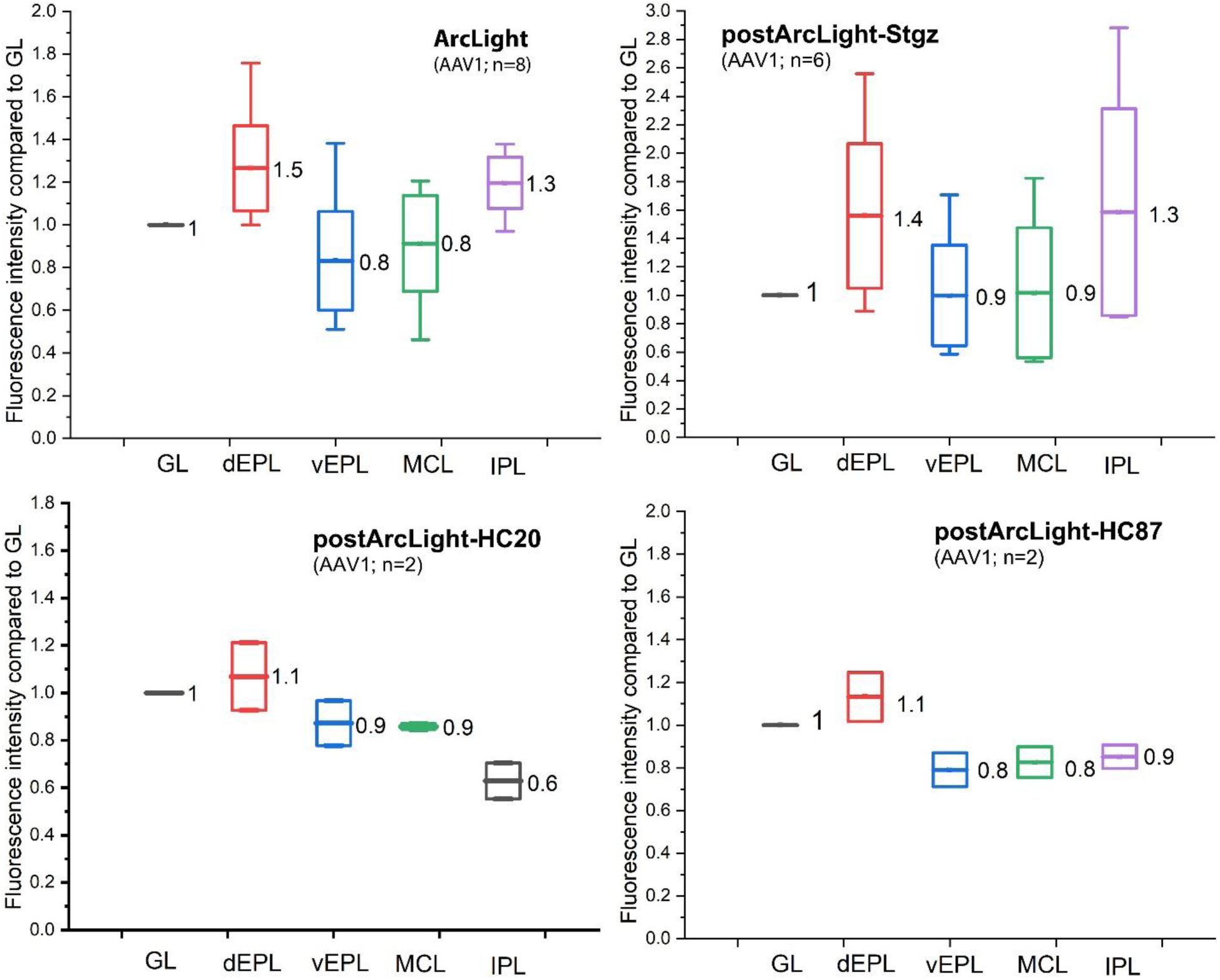
Fluorescence intensity measurements of ArcLight targeting constructs that showed non-specific subcellular expression. (A) original ArcLight; (B) postArcLight-stgz; (C) postArcLight-HC20; (D) postArcLight-HC87.

**Figure 6.**
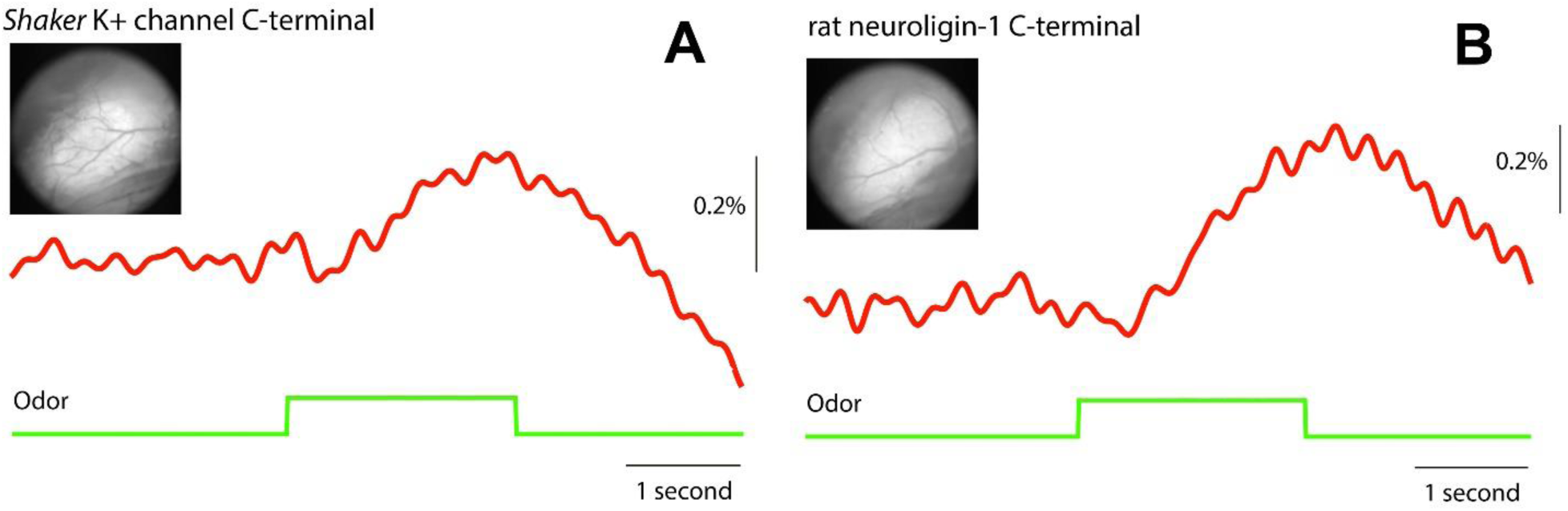
Two representatives of *in vivo* voltage fluorometric imaging on the AAV-infected fluorescent olfactory bulbs (inset) of *Tbx21-Cre* mice under odorant stimuli. (**A**) Postsynaptic targeting postArcLight-SKC; (**B**) Postsynaptic targeting postArcLight-rNL1.

#### postArcLight-HC87 (AAV1)

The postsynaptic-targeted ArcLight with anti-homer1 nanobody HC87 tag expressed approximately even in different compartment of the mitral/tufted cells (*Figure 3F, 5D*). The ArcLight expression in the dEPL was slightly higher than that in the vEPL, MCL, and IPL (*Figure 5D*).

#### postArcLight-Stgz (AAV1)

The postsynaptic-targeted ArcLight with stargazin tag showed a slightly preferential distribution within the GL and dEPL than that in vEPL, MCL, and IPL (*Figure 3G*). However, fluorescence analysis revealed no significant difference across various compartment of the mitral/tufted cells, with a larger deviation compared to other targeting constructs (*Figure 5B*). This suggests an anchoring barrier rather than subcellular trafficking issue.

The fluorescence results above suggested that the targeted ArcLight molecules reach distal axons and dendrites of mitral/tufted cells through two mechanisms: vesicular transport and lateral diffusion within the plasma membrane.

Neurexins exist and function predominantly at the presynaptic terminal. Neurexin-1β is packaged into transport vesicles and specifically directed toward the axon rather than the dendrites, and fuse to axonal membranes primarily outside of presynaptic terminals and then enter the presynaptic active zone by lateral diffusion [42, 43]. Our result with the construct preArcLight-rNxn (*Figure 3B, 4A*) was consistent with the previous observations that axonal membrane insertion of neurexin-containing vesicles was at least 2-fold higher than insertion in the somatic and dendritic membranes [42].

Neuroligin-1 is also trafficking subcellularly via transport vesicles that are carried along the dendritic microtubule network and specifically transported to the dendritic shafts and ultimately merge with the postsynaptic membrane and are dynamically exchanged at postsynaptic densities [70]. The postsynaptic targeting of neuroligin-1 can be attributed to a dendritic targeting motif and a PDZ-binding motif at its C-terminus [71]. It was suggested that the dendritic targeting motif (residues 772-803) in the C-terminus of neuroligin-1 is necessary and sufficient for dendritic targeting but not to synapse localization [72]. Our result confirmed that this dendritic targeting motif worked well for dendritic targeting (*Figure 3D*). The synaptic anchoring of neurolilgin-1 depends on the PDZ-binding interaction with PSD95, which predominantly exists at excitatory synapses, particularly at glutamatergic synapses. Most of the excitatory synapses on the olfactory neurons form directly onto the dendritic tufts in the glomeruli of the mitral and external tufted cells [73], while the lateral dendrites of mitral/tufted cells in the EPL form reciprocal synapses with granule cell dendrites for lateral inhibition [74, 75]. Our result indicated that postArcLight-rNL1 preferentially targeted to the primary dendritic tuft in the GL (*Figure 3D*), roughly 2-fold more than the lateral dendrites in the EPL (*Figure 4D*), implying that the majority of the targeted ArcLight was distributed in the excitatory synapses rather than the inhibitory synapses. Interestingly, the dendritic targeting motif (residues 772-803) of neuroligin-1 contains a gephyrin-binding motif (residues 780-794), and the Y782 phosphorylation controls the preferential binding to PSD-95 versus gephyrin [76–78]. This could be the molecular mechanism how the targeted ArcLight was unevenly distributed in both excitatory synapses in the GL dendritic tuft and the inhibitory synapses in the EPL. Moreover, it has been determined that the PDZ-binding motif of neuroligin-1 (-TTRV, residues 840-843) weakly binds to PSD95 with a K_d_ ∼120 µM [79] and the S839 phosphorylation directly preceding the PDZ-binding motif significantly reduced its PSD95 binding [80], which may be the reason why the ArcLight fluorescence was not strictly anchored in the synaptic region observed in our result (*Figure S3*). These post-translation modifications confer the capability for dynamically regulating the distribution of synaptic proteins for synaptic plasticity.

The subcellular trafficking of postArcLight-SKC exhibited a different pattern from other synaptic proteins via transport vesicle (like neurexin-1 and neuroligin-1β). It was reported that the PDZ-binding motif (-ETDV) at the *Shaker* K^+^ channel C-terminus (SKC) directs the synaptic localization of SynapCam [52], which initially distributes uniformly on the cell membrane and then clusters at the synapse. We observed a similar trafficking pattern of postArcLight-SKC that showed preferential distribution along the primary dendritic membrane extending to the dendritic tuft in the GL. Interestingly, the primary dendrites of the mitral/tufted cells illuminated by the ArcLight fluorescence, were much brighter than the lateral dendrites in the EPL (*Figure 3C*). Mitral cells typically have a single long primary dendrite extending into the glomerulus and several lateral dendrites that branch off from the primary dendrite and extend into the surrounding neuropil in the EPL. Tufted cells typically have shorter dendrites compared to mitral cells, and the dendritic tufts are more extensive extending into the EPL. The preferential trafficking along the primary dendrites instead of enrichment in the lateral dendrites could be attributed to the specific properties of the SKC targeting sequence, not just simply due to the PDZ-binding motif. Mutation of the PDZ-binding motif abolished the binding and clustering with PSD95 and SAP97 [81, 82], but it is an anchoring problem instead of a subcellular trafficking issue.

It was observed that some fluorescence of postArcLight-SKC accumulated within the soma and proximal dendrite in mitral/tufted cells, implying an issue with cell surface targeting. Overexpression of postArcLight-SKC may overwhelm the sorting and trafficking machinery, leading to mistargeting of ArcLight, including axonal mislocalization. As a result, expression level of ArcLight can influence fluorescence distribution across different layers of the main olfactory bulb (*Figure S1, upper and middle panels*). To mitigate this, we made an AAV2 virus containing the postArcLight-SKC construct to reduce ArcLight expression in the mitral/tufted cells which demonstrated preferential distribution across different layers (*Figure 5B*). However, the preferential diffusion along the primary dendritic membrane, rather than the lateral dendritic membrane, remained unchanged in different expression levels in AAV1- and AAV2-infected neurons. Additionally, the surface targeting issue persisted in AAV2-infected neurons, suggesting that this problem may be attributed to the intrinsic properties of the SKC sequence. This is not surprising, as the *Shaker* K^+^ channel C-terminal sequence contains several unique glutamine-rich regions that may originate from *Drosophila* but introduce cell surface targeting issues when expressed in mammalian neurons. We hypothesize that the postArcLight-SKC construct is expressed in the soma, transported to the somatodendritic membrane, and then diffuses along the dendritic membrane by lateral diffusion (*Figure S1, lower panel*), instead of using synaptic vesicle transport, eventually clustering in synaptic region by anchoring to the PSD. When the expression levels were high, intracellular trafficking jams occurred in the ER/Golgi pathway.

Stargazin is initially synthesized in the soma and subsequently transported to dendrites via anterograde transport vesicles, maintaining proper cell membrane targeting. Stargazin undergoes dynamic phosphorylation and dephosphorylation at its C-terminus, modulating its localization in and out of the postsynaptic region. This phosphorylation process regulates the postsynaptic localization of AMPA receptors [83]. In the postArcLight-Stgz targeting construct, three phosphomimetic mutations (S239D, S240D, and S253D) were introduced to detach the C-terminus from the cell membrane, promoting stargazin movement to the postsynaptic density (PSD). However, T321 within the PDZ-binding motif (-TTPV, residues 320-323) is surrounded by consensus sequences for phosphorylation by PKA, PKC, CaMKII, and MAPKs. Phosphomimetic mutations of T321E or T321D eliminated coclustering of stargazin and PSD-95 [56]. Consequently, T321 phosphorylation of the stargazin PDZ-binding motif can significantly attenuate anchoring to PSD-95, consistent with our observations (*Figure 3G, 5B*).

Homer1 is a postsynaptic scaffolding protein in excitatory synapses, confined to the synaptic region without extending into the perisynaptic zone [62]. The Homer1 layer is situated approximately 80 nm from the synaptic cleft [63, 84]. However, the cytoplasmic part of postArcLight-HC20/HC87 is less than 20 nm in total length, including the superecliptic pHluorin (∼4.2 nm), and the 15-residue linker (∼5 nm) [85] between ArcLight and the Homer1 nanobody (HC20/HC87). Our experiments showed that the fluorescence of postArcLight-HC20/HC87 was distributed not only in the primary dendrites of the GL and lateral dendrites in the EPL, but also more evenly throughout the mitral/tufted cells (*Figure 3E, 3F, 5C and 5D*). This broader distribution likely results from the absence of dendritic sorting/trafficking sequences and the construct’s inability to reach the Homer1 layer.

It is worth noting that the expression level of the ArcLight targeting construct may affect the fluorescence distribution in the different layers of olfactory bulb depending on the subcellular sorting/trafficking pattern. It was observed that the ArcLight fluorescence in some neurons with postArcLight-SKC overexpression stocked in the inner membrane of the soma projected along the primary dendrite (*Supplemental material, Figure S1*), which implied a certain difficulty for cell membrane targeting with the SKC tag. We made AAV2 virus with the postArcLight-SKC construct to compare with AAV1 virus as AAV2 virus has less expression than AAV1 in the mitral/tufted cells. The AAV2 of postArcLight-SKC showed a preferential distribution (*Figure 3C, 4C*) while the AAV1 version was more evenly expressed in different layers than the AAV2 version (*Figure S1*) which may be due to the high expression level. However, the trafficking pattern of overexpressed postArcLight-SKC seemed similar between the AAV1 and AAV2 serotypes with preferential distribution in the primary dendritic tuft along the long projection from the exceptional high expression in the soma. Neither the AAV1 nor the AAV2 of postArcLight-HC20 showed a preferential distribution (*Figure S2*).

High magnification images revealed both membrane expression and punctate expression in the glomerular dendritic tuft and the primary dendrites of the mitral/tufted cells with all of constructs (*Figure S3*). We hypothesize that the punctate fluorescence pattern may be due to the ArcLight clusters around postsynaptic structures while the ArcLight also distributed along the dendritic membrane suggesting that the ArcLight may not be exclusively anchored in the synaptic region. Similarly, there were also fluorescent puncta on the axon shafts in the IPL with the preArcLight-rNxn (*Figure S4*). The axons of tufted cell can form axodendritic synapses with proximal dendrites of granule cells in the IPL [86]. The puncta on the axon membrane in the IPL implied presynaptic density of the axodendritic synapses.

### *In Vivo* Voltage Imaging

The *in vivo* voltage fluorometric imaging on the AAV-injected olfactory bulbs of *Tbx21-Cre* mice showed that the subcellularly targeted ArcLight had a similar fractional fluorescence change as the control ArcLight (*Table 1*). It was confirmed that the targeted Arclight can be functional for voltage sensitivity in the live animal under odorant stimuli. In this in vivo voltage imaging experiments, we cannot distinguish single glomerulus or single neuron with low-magnification microscopy. Further investigation with multiple photon microscopy would be helpful to distinguish the voltage signals from different compartments of mitral/tufted cells in mouse olfactory bulb.

**Table 1.** Voltage response in the AAV-infected olfactory bulbs of *Tbx21-Cre* mice.

| CONSTRUCTS | AAV SEROTYPE | $\Delta F/F$ (%) | MICE* |
| --- | --- | --- | --- |
| Control ArcLight | AAV1 | $0.37 \pm 0.05$ | 3 |
| preArcLight-rNxn | AAV1 | $0.32 \pm 0.07$ | 2 |
| somaArcLight-KvLkTn | AAV2 | $0.46 \pm 0.10$ | 7 |
| postArcLight-rNL1 | AAV2 | $0.48 \pm 0.12$ | 4 |
| postArcLight-SKC | AAV1 | $0.45 \pm 0.06$ | 5 |
| | AAV2 | $0.18 \pm 0.04$ | 3 |
| postArcLight-Stgz | AAV1 | $0.34 \pm 0.09$ | 3 |
| postArcLight-HC20 | AAV1 | $0.37 \pm 0.09$ | 3 |
| | AAV2 | $0.47 \pm 0.13$ | 4 |
| postArcLight-HC87 | AAV1 | $0.22 \pm 0.07$ | 2 |

\* The numbers of *Tbx21-Cre* mice used in the *in vivo* voltage imaging experiment.

## DISCUSSION

In this study, we verified the effectiveness of targeting ArcLight constructs that can be specifically expressed and targeted in different subcellular compartments in the mitral/tufted cells of the olfactory bulb in *Tbx21-Cre* mice. The targeting sequences used in the targeting ArcLight constructs engage distinct trafficking machinery and exhibit varying binding affinities for anchoring target proteins (e.g. PSD-95, Homer-1, etc.), leading to diverse subcellular distributions and trafficking patterns. The ArcLight molecules can travel to distal axon/dendrites in the mitral/tufted cells by either transport vesicles or by lateral diffusion in the plasma membrane, depending on the targeting sequences.

The *in vivo* targeted ArcLight expression in the mitral/tufted cells of mouse olfactory bulb was achieved with different targeting sequences. The different compartmental targeting includes specific sorting/trafficking machinery and the anchoring interactions with synaptic scaffolding proteins, which are modulated by post-translation modification (e.g. phosphorylation) and also affected by the protein sequence context in the targeting constructs. In this study, we confirmed that the targeted ArcLight molecules were still functional for voltage sensitivity in the live animal under odorant stimuli. However, we cannot distinguish the voltage signal from different compartments, like dendritic or axonal signals, with 1P wide-field microscopy that cannot provide high spatial resolution in Z and XY axis. The odorant response in the olfactory bulb is complicated with regional differences and not every glomerulus gets activated and not every mitral/tufted cell started firing under odorant stimuli, and furthermore intrinsic inhibition would function afterwards. The cross-talk of dendritic integration [87, 88] and axon computation [89] in the mitral/tufted cell and granule cells in olfactory bulb may be much more complicated than a simple signal transmission [90]. The SNR limit may be also related to the spatiotemporal resolution of the low-magnification microscopy that we used in the *in vivo* voltage imaging.

With the development of multiphoton microscopy and superresolution microscopy, the synaptic-targeting expression of GEVIs/GECIs in different functional compartment of neuron can be applied to further research in mental diseases involving synaptic disorders.

## Abbreviations

PSD: postsynaptic density
ONL: Olfactory Neuron Layer
GL: Glomerular Layer
EPL: External Plexiform Layer
dEPL: dorsal EPL
vEPL: ventral EPL
MCL: Mitral Cell Layer
IPL: inner Plexiform Layer
GCL: Granule Cell Layer
rNxn: rat Neurexin-1β C-terminal tag
KvLkTn: Kv2.1-linker-Tlcn C-terminal tag
SKC: *Shaker* K^+^ channel C-terminal tag
rNL1: rat Neuroligin-1 C-terminal tag
Stgz: stargazin C-terminal with phosphomimetic charge mutations
HC20: HC20 anti-homer1 nanobody tag
HC87: HC87 anti-homer1 nanobody tag.

## Data Availability

The data that support the findings of this study are available from the authors.

## Author Contribution

S. Z. designed and made the ArcLight targeting constructs in this study, performed the experiments of molecular biology, electrophysiology, confocal imaging, and AAV injection into mouse olfactory bulb. S.Z. and L.B.C. did the *in vivo* voltage imaging, analyzed the experimental data and wrote the paper.

## Acknowledgements

In heartfelt acknowledgment, I wish to extend my deepest gratitude to my mentor, the late Dr. Lawrence (Larry) Baruch Cohen, who suddenly passed away in April of 2023. Larry was an illustrious pioneer in the development of optical methodologies for monitoring membrane potential in alive cells, especially action potential in neurons. Remarkably, Larry sustained an active laboratory for over 50 years, never ceasing his involvement in experiments and data analysis as well as writing grants, including this paper, until sudden passing. Larry’s sharp intellect, analytical acumen, and thoughtful nature were legendary, often concealed by his deliberate speech. Larry was more than an outstanding scientist, but also an exceptional mentor. In times of funding uncertainty, Larry unflinchingly dipped into his own pocket to support his students and postdocs, exemplifying his commitment to the pursuit of scientific excellence. Larry always wore a red shirt faithfully for at least 40 years as his portrait symbol. Larry enjoyed simple pleasures: fine wine and friendly one-dollar bets. The wall in front of his office desk was adorned with many one-dollar bills, tokens of his playful victories, some from me. The last bet between Larry and I was $100 for a scientific argument, but unfortunately, we did not finish it. As the last member in Larry’s lab at Yale university, I had benefited a lot from Larry, and I pledge to continue the marathon he started, running towards the scientific goals.

Here I am grateful to my postdoctoral committee members at Yale University - Dr. Michael Caplan, Dr. Biff Forbush, and Dr. David Zenisek - for their kind guidance. I would also like to thank Dr. William N. Ross (New York Medical College), Dr. Brian Salzberg (University of Pennsylvania), Dr. Dejan Zecevic (Yale University), Dr. Thomas C. Südhof (Stanford University), Dr. Ehud Y. Isacoff (UC Berkeley), Dr. Bradley Baker (Korea Institute of Science and Technology), Dr. Douglas Storace (Florida State University), and Dr. Xiaowei Hou (University of Cincinnati) for their critical feedback and valuable suggestions on this paper and our future research directions.

The SynView plasmids [29] were a kind gift from Dr. Thomas C. Südhof at Stanford University. The SynapGCaMP plasmid [51] for *Shaker* K^+^ Channel C-terminal sequence was a kind gift from Dr. Ehud Y. Isacoff at UC Berkeley. The optimized ArcLight plasmid (Addgene #100038) was a gift from Dr. Vincent Pieribone at Yale university & John B. Pierce Laboratory. The plasmids of anti-Homer1 nanobody HC20 & HC87 (Addgene plasmid #135220 & #135223) [64] were a gift from Dr. James Trimmer at UC Davis. The plasmid of pAAV-EF1α-FRT-FLEX-GtACR2-EYFP-Kv2.1C-linker-TlcnC (Addgene plasmid #114377) [48] was a gift from Dr. Mingshan Xue. The plasmid of Stargazin-GFP-LOVpep (Addgene plasmid # 80406) [60] was a gift from Dr. Michael Glotzer.

## Supplemental Data

**Figure S1.**
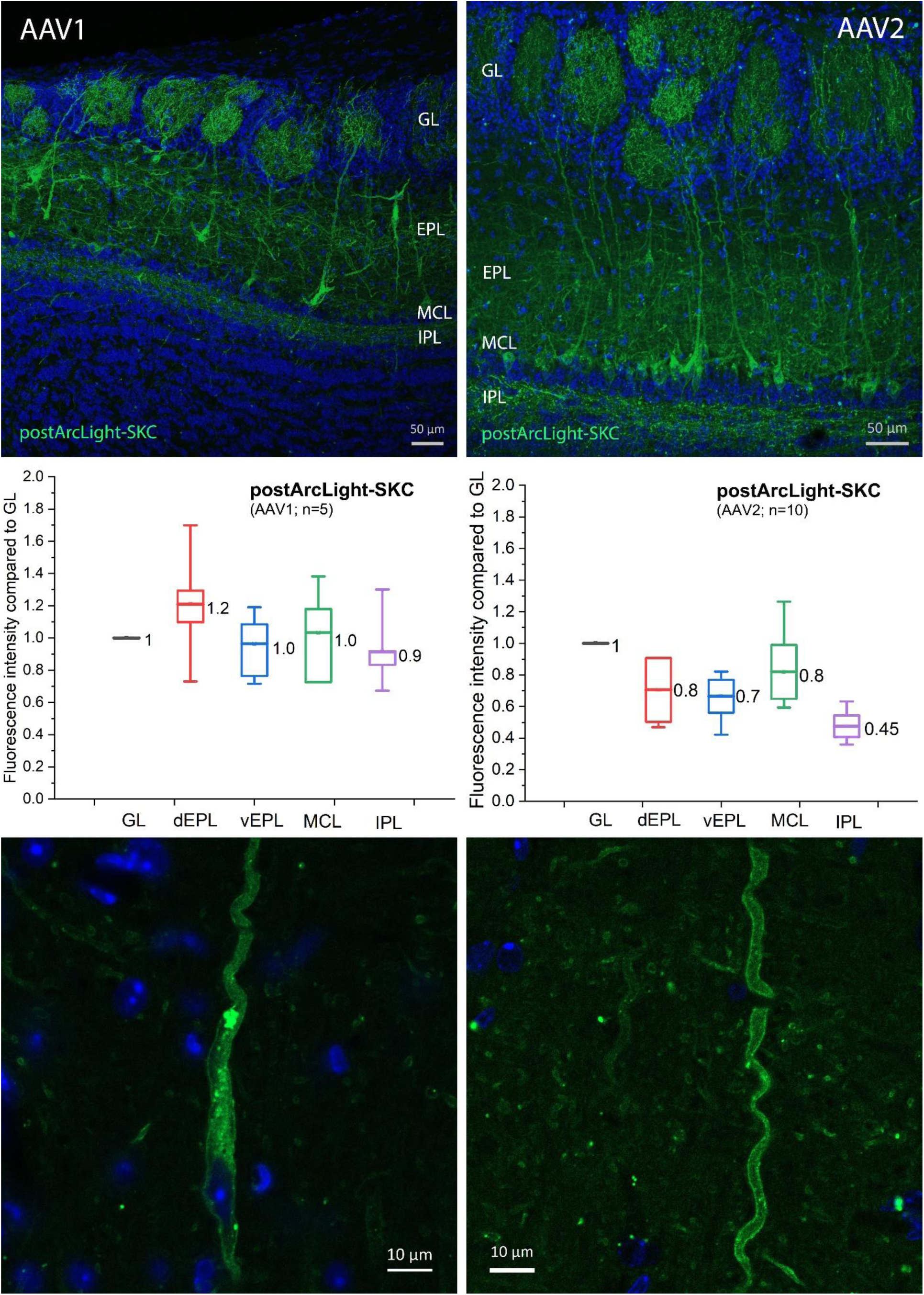
Different AAV serotypes of postArcLight-SKC. The expression level of AAV1 virus is generally higher than AAV2 virus in the olfactory bulb of Tbx21-Cre mice. (**Upper Panel**) While the expression level affected the fluorescence distribution in the different layers of mouse olfactory bulb, the trafficking pattern of postArcLight-SKC seemed similar between the AAV1 and AAV2 serotypes with preferential distribution in the primary dendritic tuft along the long projection from the high expression in the soma. (**Middle panel**) The statistical comparison between the AAV1 and AAV2 serotypes of postArcLight-SKC. (**Lower panel**) High magnification images showed that the targeted ArcLight in the soma (left) and dendritic membrane, respectively, while some of them were accumulated in the inner membrane with the preferential dendritic side. This tendency was also observed in the soma at the MCL in the upper panel.

**Figure S2.**
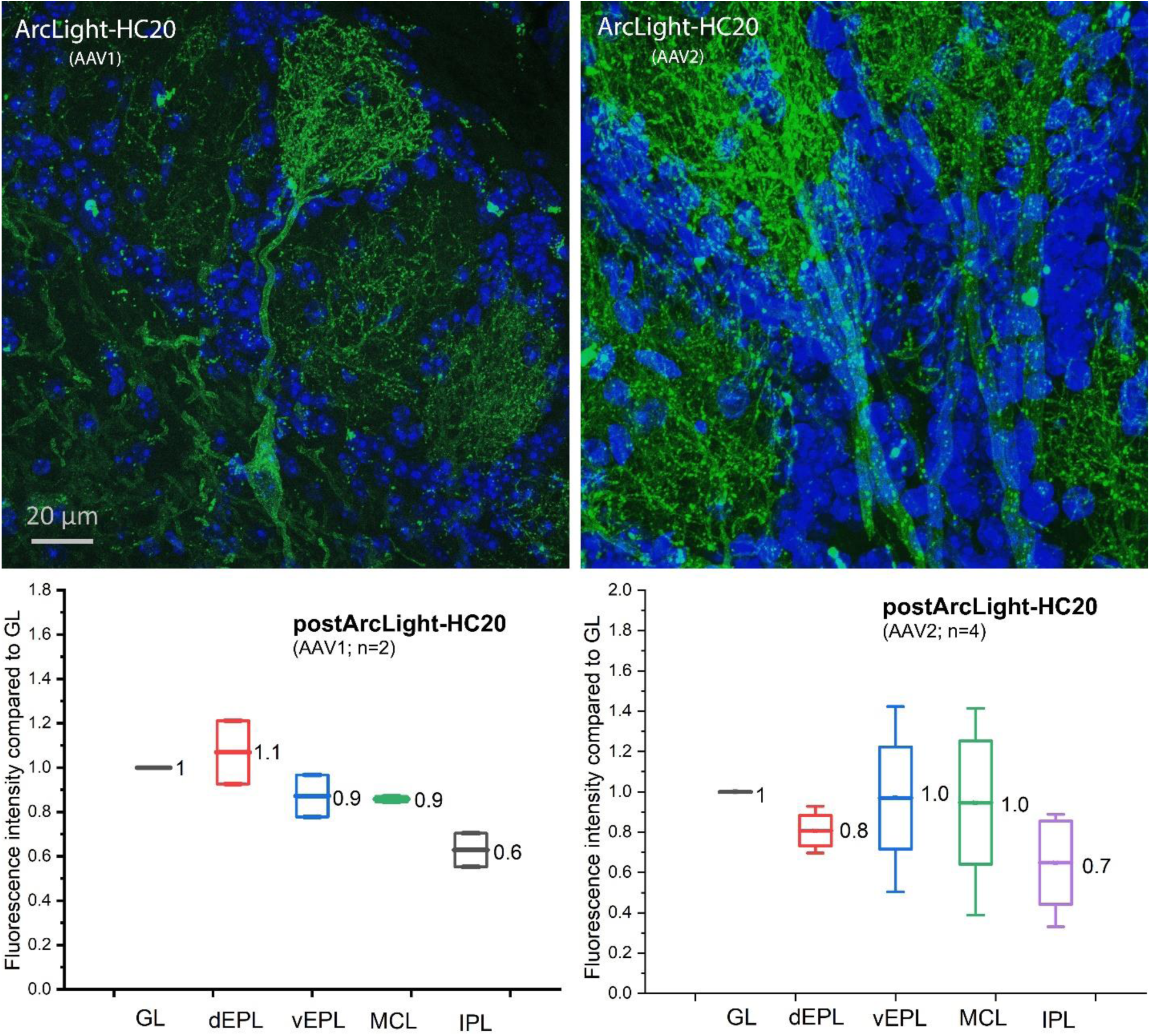
Different AAV serotypes of postArcLight-HC20. The expression level of AAV1 and AAV2 were different, but both AAV viruses made with postArcLight-HC20 did not show different distribution pattern in the compartments of the mitral/tufted cells. (**Upper Left**) An external tufted cell was shown as the primary dendrite along to the dendritic tuft in the glomerulus. The spotty dots on the dendritic membrane may be the postsynaptic density where the anti-Homer1 nanobody HC20 interacts with the scaffolding protein Homer1.

**Figure S3.**
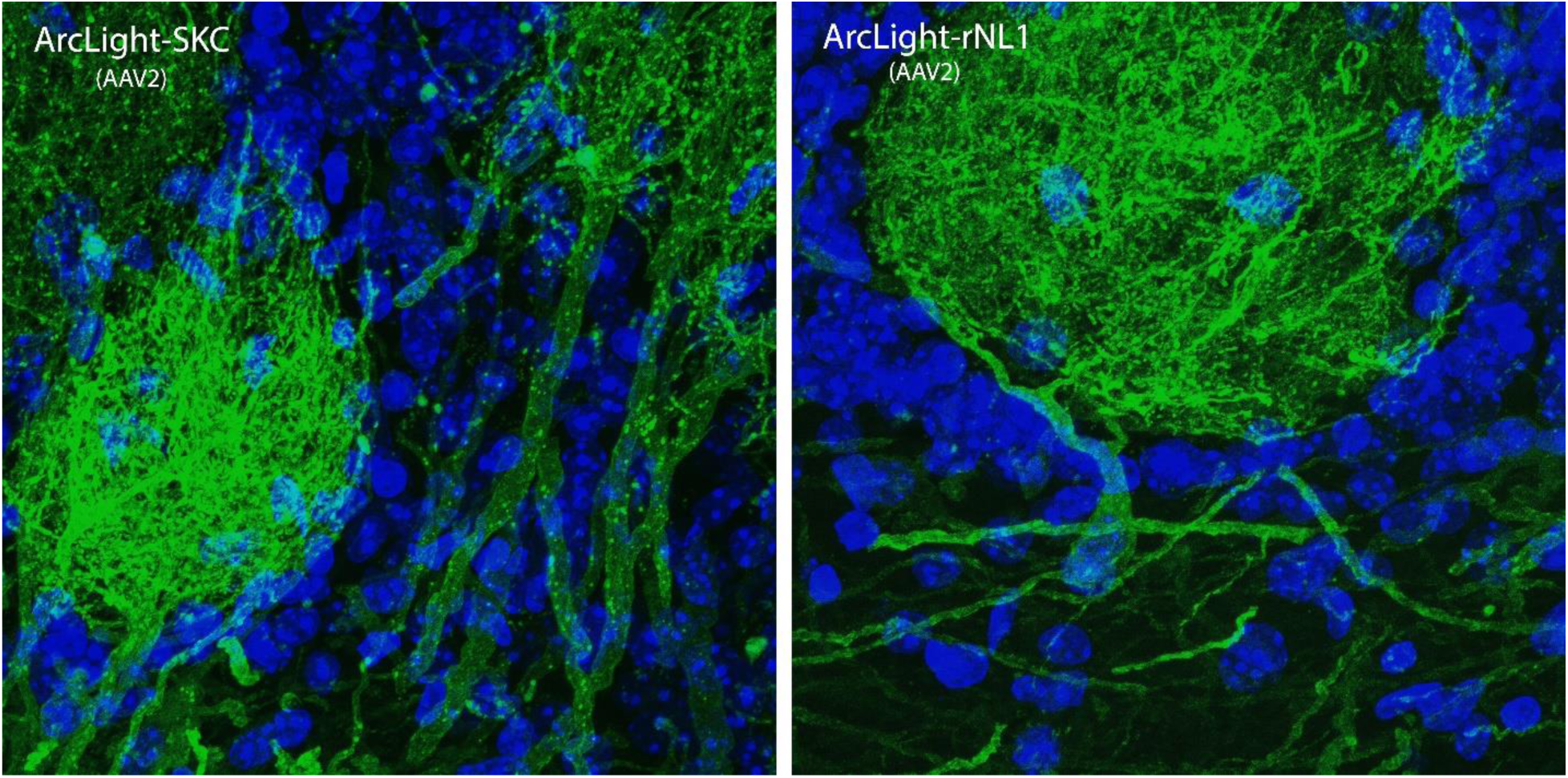
Postsynaptic targeting ArcLight shown enriched accumulation (bright spots) on the distal dendrites in the GL as well as the lateral diffusion along the dendrite membrane.

**Figure S4.**
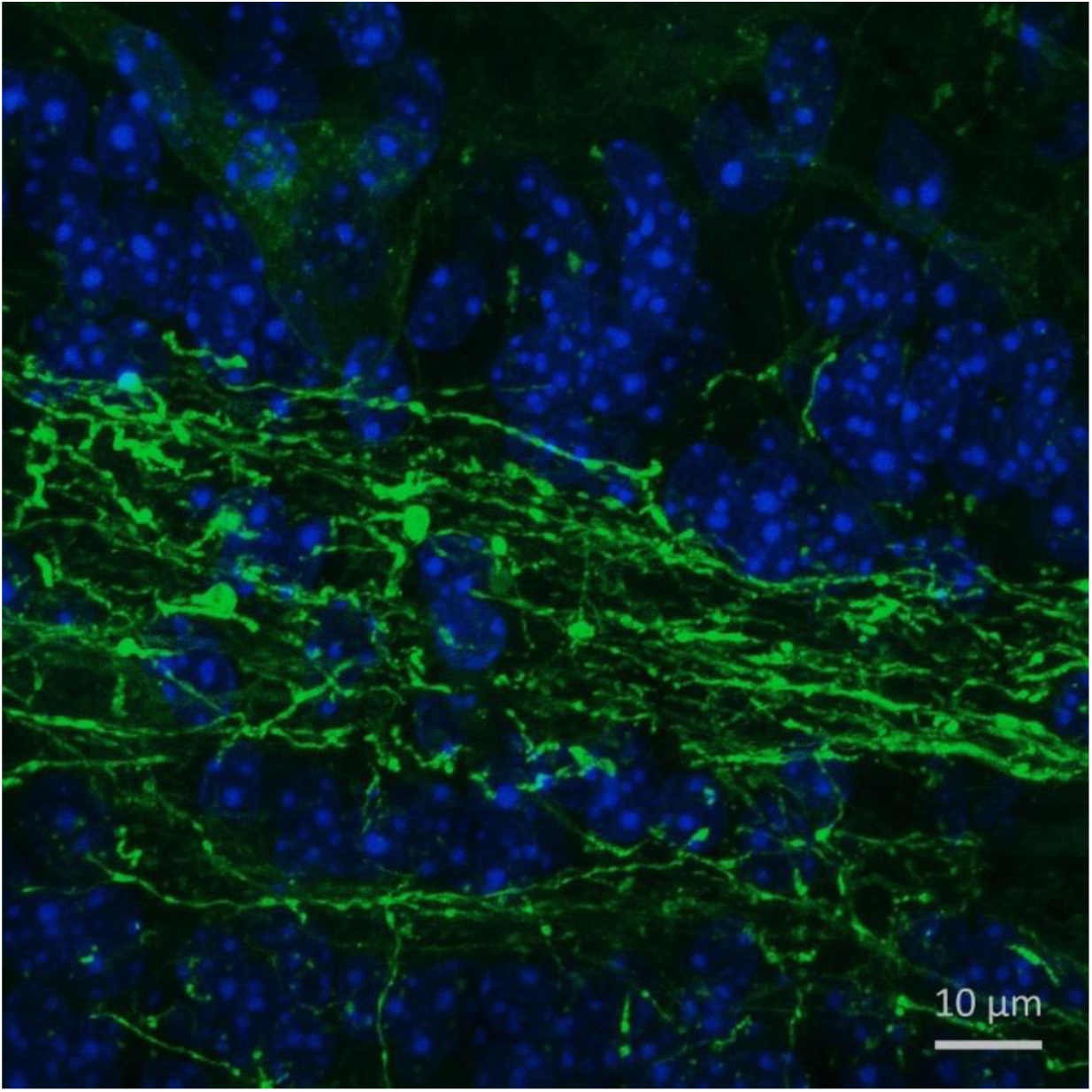
Presynaptic targeting ArcLight shown enriched accumulation (bright spots).

## Uncategorized References

1. Sanchez, C., et al., Detection of Ca2+ transients near ryanodine receptors by targeting fluorescent Ca2+ sensors to the triad. J Gen Physiol, 2021. 153(4).

2. Zhu, M.H., et al., Population imaging discrepancies between a genetically-encoded calcium indicator (GECI) versus a genetically-encoded voltage indicator (GEVI). Scientific Reports, 2021. 11(1).

3. Storace, D., et al., Genetically Encoded Protein Sensors of Membrane Potential. Adv Exp Med Biol, 2015. 859: p. 493–509.

4. Nguyen, C., et al., Simultaneous voltage and calcium imaging and optogenetic stimulation with high sensitivity and a wide field of view. Biomed Opt Express, 2019. 10(2): p. 789–806.

5. Sepehri Rad, M., et al., Voltage and Calcium Imaging of Brain Activity. Biophys J, 2017. 113(10): p. 2160–2167.

6. Platisa, J., et al., Voltage imaging in the olfactory bulb using transgenic mouse lines expressing the genetically encoded voltage indicator ArcLight. Sci Rep, 2022. 12: p. 1875.

7. Leong, L.M. and D.A. Storace, Imaging different cell populations in the mouse olfactory bulb using the genetically encoded voltage indicator ArcLight. Neurophotonics, 2024. 11(3).

8. Zang, Y. and E. Marder, Neuronal morphology enhances robustness to perturbations of channel densities. Proc Natl Acad Sci U S A, 2023. 120(8): p. e2219049120.

9. Francioni, V. and M.T. Harnett, Rethinking Single Neuron Electrical Compartmentalization: Dendritic Contributions to Network Computation. Neuroscience, 2022. 489: p. 185–199.

10. Stuart, G.J. and N. Spruston, Dendritic integration: 60 years of progress. Nature Neuroscience, 2015. 18(12): p. 1713–1721.

11. Davila, H.V., et al., Large Change in Axon Fluorescence That Provides a Promising Method for Measuring Membrane-Potential. Nature-New Biology, 1973. 241(109): p. 159–160.

12. Salzberg, B.M., H.V. Davila, and L.B. Cohen, Optical Recording of Impulses in Individual Neurons of an Invertebrate Central Nervous-System. Nature, 1973. 246(5434): p. 508–509.

13. Landau, A.T., et al., Dendritic branch structure compartmentalizes voltage-dependent calcium influx in cortical layer 2/3 pyramidal cells. Elife, 2022. 11.

14. Armbruster, M., et al., Neuronal activity drives pathway-specific depolarization of peripheral astrocyte processes. Nat Neurosci, 2022. 25(5): p. 607–616.

15. Short, S.M., et al., The stochastic nature of action potential backpropagation in apical tuft dendrites. Journal of Neurophysiology, 2017. 118(2): p. 1394–1414.

16. Ozbay, B.N., et al., Three dimensional two-photon brain imaging in freely moving mice using a miniature fiber coupled microscope with active axial-scanning. Sci Rep, 2018. 8(1): p. 8108.

17. Platisa, J., et al., High-speed low-light in vivo two-photon voltage imaging of large neuronal populations. Nat Methods, 2023. 20: p. 1095–1103.

18. Wang, T., et al., Three-photon imaging of mouse brain structure and function through the intact skull. Nat Methods, 2018. 15: p. 789–792.

19. Streich, L., et al., High-resolution structural and functional deep brain imaging using adaptive optics three-photon microscopy. Nat Methods, 2021. 18: p. 1253–1258.

20. Lecoq, J.A., R. Boehringer, and B.F. Grewe, Deep brain imaging on the move. Nat Methods, 2023. 20(4): p. 495–496.

21. Ishii, H., et al., Focusing new light on brain functions: multiphoton microscopy for deep and super-resolution imaging. Neuroscience Research, 2022. 179: p. 24–30.

22. Lim, S.T., et al., A novel targeting signal for proximal clustering of the Kv2.1 K+ channel in hippocampal neurons. Neuron, 2000. 25(2): p. 385–97.

23. Abdelfattah, A.S., et al., Bright and photostable chemigenetic indicators for extended in vivo voltage imaging. Science, 2019. 365(6454): p. 699-+.

24. Adam, Y., et al., Voltage imaging and optogenetics reveal behaviour-dependent changes in hippocampal dynamics. Nature, 2019. 569(7756): p. 413-+.

25. Villette, V., et al., Ultrafast Two-Photon Imaging of a High-Gain Voltage Indicator in Awake Behaving Mice. Cell, 2019. 179(7): p. 1590–1608 e23.

26. Mitsui, S., et al., A novel phenylalanine-based targeting signal directs telencephalin to neuronal dendrites. J Neurosci, 2005. 25(5): p. 1122–31.

27. Guerrero, G., et al., Heterogeneity in synaptic transmission along a Drosophila larval motor axon. Nat Neurosci, 2005. 8(9): p. 1188–96.

28. Kim, J., et al., mGRASP enables mapping mammalian synaptic connectivity with light microscopy. Nat Methods, 2011. 9(1): p. 96–102.

29. Tsetsenis, T., et al., Direct visualization of trans-synaptic neurexin-neuroligin interactions during synapse formation. J Neurosci, 2014. 34(45): p. 15083–96.

30. Jin, L., et al., Single action potentials and subthreshold electrical events imaged in neurons with a fluorescent protein voltage probe. Neuron, 2012. 75(5): p. 779–85.

31. Bando, Y., et al., Comparative Evaluation of Genetically Encoded Voltage Indicators. Cell Rep, 2019. 26(3): p. 802–813 e4.

32. Milosevic, M.M., et al., In Vitro Testing of Voltage Indicators: Archon1, ArcLightD, ASAP1, ASAP2s, ASAP3b, Bongwoori-Pos6, BeRST1, FlicR1, and Chi-VSFP-Butterfly. eNeuro, 2020. 7(5).

33. Lee, S., et al., A trafficking motif alters GEVI activity implicating persistent protein interactions at the membrane. Biophysical Reports, 2022. 2(2).

34. Stockklausner, C., et al., A sequence motif responsible for ER export and surface expression of Kir2.0 inward rectifier K(+) channels. FEBS Lett, 2001. 493(2-3): p. 129–33.

35. Ma, D., et al., Role of ER export signals in controlling surface potassium channel numbers. Science, 2001. 291(5502): p. 316–9.

36. Stockklausner, C. and N. Klocker, Surface expression of inward rectifier potassium channels is controlled by selective Golgi export. J Biol Chem, 2003. 278(19): p. 17000–5.

37. Hofherr, A., B. Fakler, and N. Klocker, Selective Golgi export of Kir2.1 controls the stoichiometry of functional Kir2.x channel heteromers. J Cell Sci, 2005. 118(Pt 9): p. 1935–43.

38. Kwon, T., et al., Attenuation of Synaptic Potentials in Dendritic Spines. Cell Rep, 2017. 20(5): p. 1100–1110.

39. Platisa, J., et al., Directed Evolution of Key Residues in Fluorescent Protein Inverses the Polarity of Voltage Sensitivity in the Genetically Encoded Indicator ArcLight. ACS Chem Neurosci, 2017. 8(3): p. 513–523.

40. Dean, C. and T. Dresbach, Neuroligins and neurexins: linking cell adhesion, synapse formation and cognitive function. Trends Neurosci, 2006. 29(1): p. 21–9.

41. Sudhof, T.C., Neuroligins and neurexins link synaptic function to cognitive disease. Nature, 2008. 455(7215): p. 903–11.

42. Fairless, R., et al., Polarized targeting of neurexins to synapses is regulated by their C-terminal sequences. J Neurosci, 2008. 28(48): p. 12969–81.

43. Neupert, C., et al., Regulated Dynamic Trafficking of Neurexins Inside and Outside of Synaptic Terminals. J Neurosci, 2015. 35(40): p. 13629–47.

44. Trimmer, J.S., Immunological identification and characterization of a delayed rectifier K+ channel polypeptide in rat brain. Proc Natl Acad Sci U S A, 1991. 88(23): p. 10764–8.

45. Jensen, C.S., et al., Specific sorting and post-Golgi trafficking of dendritic potassium channels in living neurons. J Biol Chem, 2014. 289(15): p. 10566–10581.

46. Jensen, C.S., et al., Trafficking of Kv2.1 Channels to the Axon Initial Segment by a Novel Nonconventional Secretory Pathway. J Neurosci, 2017. 37(48): p. 11523–11536.

47. Wu, C., et al., rAAV-mediated subcellular targeting of optogenetic tools in retinal ganglion cells in vivo. PLoS One, 2013. 8(6): p. e66332.

48. Messier, J.E., et al., Targeting light-gated chloride channels to neuronal somatodendritic domain reduces their excitatory effect in the axon. Elife, 2018. 7: p. e38506.

49. Song, J.Y., et al., Neuroligin 1 is a postsynaptic cell-adhesion molecule of excitatory synapses. Proceedings of the National Academy of Sciences of the United States of America, 1999. 96(3): p. 1100–1105.

50. Shipman, S.L., et al., Functional dependence of neuroligin on a new non-PDZ intracellular domain. Nature Neuroscience, 2011. 14(6): p. 718–U388.

51. Newman, Z.L., et al., Input-Specific Plasticity and Homeostasis at the Drosophila Larval Neuromuscular Junction. Neuron, 2017. 93(6): p. 1388–1404 e10.

52. Zito, K., et al., Synaptic clustering of Fascilin II and Shaker: essential targeting sequences and role of Dlg. Neuron, 1997. 19(5): p. 1007–16.

53. Yamazaki, M., et al., A novel action of stargazin as an enhancer of AMPA receptor activity. Neurosci Res, 2004. 50(4): p. 369–74.

54. Tomita, S., et al., Bidirectional synaptic plasticity regulated by phosphorylation of stargazin-like TARPs. Neuron, 2005. 45(2): p. 269–77.

55. Choi, J., et al., Phosphorylation of stargazin by protein kinase A regulates its interaction with PSD-95. J Biol Chem, 2002. 277(14): p. 12359–63.

56. Stein, E.L. and D.M. Chetkovich, Regulation of stargazin synaptic trafficking by C-terminal PDZ ligand phosphorylation in bidirectional synaptic plasticity. J Neurochem, 2010. 113(1): p. 42–53.

57. Opazo, P., et al., CaMKII Triggers the Diffusional Trapping of Surface AMPARs through Phosphorylation of Stargazin. Neuron, 2010. 67(2): p. 239–252.

58. Sumioka, A., D. Yan, and S. Tomita, TARP phosphorylation regulates synaptic AMPA receptors through lipid bilayers. Neuron, 2010. 66(5): p. 755–67.

59. Bats, C., L. Groc, and D. Choquet, The interaction between Stargazin and PSD-95 regulates AMPA receptor surface trafficking. Neuron, 2007. 53(5): p. 719–34.

60. Wagner, E. and M. Glotzer, Local RhoA activation induces cytokinetic furrows independent of spindle position and cell cycle stage. J Cell Biol, 2016. 213(6): p. 641–9.

61. Hafner, A.S., et al., Lengthening of the Stargazin Cytoplasmic Tail Increases Synaptic Transmission by Promoting Interaction to Deeper Domains of PSD-95. Neuron, 2015. 86(2): p. 475–89.

62. Tao-Cheng, J.H., et al., Homer is concentrated at the postsynaptic density and does not redistribute after acute synaptic stimulation. Neuroscience, 2014. 266: p. 80–90.

63. Gutierrez-Mecinas, M., et al., Immunostaining for Homer reveals the majority of excitatory synapses in laminae I-III of the mouse spinal dorsal horn. Neuroscience, 2016. 329: p. 171–81.

64. Dong, J.X., et al., A toolbox of nanobodies developed and validated for use as intrabodies and nanoscale immunolabels in mammalian brain neurons. Elife, 2019. 8.

65. Brandstatter, J.H., O. Dick, and T.M. Boeckers, The postsynaptic scaffold proteins ProSAP1/Shank2 and Homer1 are associated with glutamate receptor complexes at rat retinal synapses. J Comp Neurol, 2004. 475(4): p. 551–63.

66. Mitsui, S., et al., Genetic visualization of the secondary olfactory pathway in Tbx21 transgenic mice. Neural Syst Circuits, 2011. 1(1): p. 5.

67. Storace, D.A. and L.B. Cohen, Measuring the olfactory bulb input-output transformation reveals a contribution to the perception of odorant concentration invariance. Nat Commun, 2017. 8(1): p. 81.

68. Vucinic, D., L.B. Cohen, and E.K. Kosmidis, Interglomerular center-surround inhibition shapes odorant-evoked input to the mouse olfactory bulb in vivo. J Neurophysiol, 2006. 95(3): p. 1881–7.

69. Imamura, F., A. Ito, and B.J. LaFever, Subpopulations of Projection Neurons in the Olfactory Bulb. Front Neural Circuits, 2020. 14: p. 561822.

70. Schapitz, I.U., et al., Neuroligin 1 Is Dynamically Exchanged at Postsynaptic Sites. Journal of Neuroscience, 2010. 30(38): p. 12733–12744.

71. Dresbach, T., et al., Synaptic targeting of neuroligin is independent of neurexin and SAP90/PSD95 binding. Mol Cell Neurosci, 2004. 27(3): p. 227–35.

72. Rosales, C.R., et al., A cytoplasmic motif targets neuroligin-1 exclusively to dendrites of cultured hippocampal neurons. Eur J Neurosci, 2005. 22(9): p. 2381–6.

73. Bourne, J.N. and N.E. Schoppa, Three-dimensional synaptic analyses of mitral cell and external tufted cell dendrites in rat olfactory bulb glomeruli. J Comp Neurol, 2017. 525(3): p. 592–609.

74. Aghvami, S.S., Y. Kubota, and V. Egger, Anatomical and Functional Connectivity at the Dendrodendritic Reciprocal Mitral Cell-Granule Cell Synapse: Impact on Recurrent and Lateral Inhibition. Frontiers in Neural Circuits, 2022. 16.

75. Burton, S.D., C.M. Malyshko, and N.N. Urban, Fast-spiking interneuron detonation drives high-fidelity inhibition in the olfactory bulb. bioRxiv, 2024.

76. Giannone, G., et al., Neurexin-1 beta Binding to Neuroligin-1 Triggers the Preferential Recruitment of PSD-95 versus Gephyrin through Tyrosine Phosphorylation of Neuroligin-1. Cell Reports, 2013. 3(6): p. 1996–2007.

77. Bemben, M.A., et al., CaMKII phosphorylation of neuroligin-1 regulates excitatory synapses. Nat Neurosci, 2014. 17(1): p. 56–64.

78. Letellier, M., et al., Optogenetic control of excitatory post-synaptic differentiation through neuroligin-1 tyrosine phosphorylation. Elife, 2020. 9.

79. Irie, M., et al., Binding of neuroligins to PSD-95. Science, 1997. 277(5331): p. 1511–5.

80. Jeong, J., et al., PSD-95 binding dynamically regulates NLGN1 trafficking and function. Proc Natl Acad Sci U S A, 2019. 116(24): p. 12035–12044.

81. Kim, E. and M. Sheng, Differential K+ channel clustering activity of PSD-95 and SAP97, two related membrane-associated putative guanylate kinases. Neuropharmacology, 1996. 35(7): p. 993–1000.

82. Magidovich, E., et al., Intrinsic disorder in the C-terminal domain of the Shaker voltage-activated K+ channel modulates its interaction with scaffold proteins. Proc Natl Acad Sci U S A, 2007. 104(32): p. 13022–7.

83. Matsuda, S., et al., Stargazin regulates AMPA receptor trafficking through adaptor protein complexes during long-term depression. Nat Commun, 2013. 4: p. 2759.

84. Dani, A., et al., Superresolution imaging of chemical synapses in the brain. Neuron, 2010. 68(5): p. 843–56.

85. Hofmann, H., et al., Polymer scaling laws of unfolded and intrinsically disordered proteins quantified with single-molecule spectroscopy. Proc Natl Acad Sci U S A, 2012. 109(40): p. 16155–60.

86. Sun, X., et al., CCKergic Tufted Cells Differentially Drive Two Anatomically Segregated Inhibitory Circuits in the Mouse Olfactory Bulb. J Neurosci, 2020. 40(32): p. 6189–6206.

87. Corlukic, M., J. Krpan, and M. Stojanovic, Dendritic Integration Theory as a Cellular Bridge for the Current Major Consciousness Theories. Psihologijske Teme, 2023. 32(3): p. 529–554.

88. Mueller, M. and V. Egger, Dendritic integration in olfactory bulb granule cells upon simultaneous multispine activation: Low thresholds for nonlocal spiking activity. Plos Biology, 2020. 18(9).

89. Alcami, P. and A. El Hed, Axonal Computations. Frontiers in Cellular Neuroscience, 2019. 13: p. 413.

90. Goaillard, J.M., et al., Diversity of Axonal and Dendritic Contributions to Neuronal Output. Frontiers in Cellular Neuroscience, 2020. 13.

